# Country-wide genomic surveillance of SARS-CoV-2 strains

**DOI:** 10.1101/2021.06.08.447365

**Authors:** Kishan Kalia, Gayatri Saberwal, Gaurav Sharma

## Abstract

Genomic surveillance has enabled the identification of several SARS-CoV-2 variants, allowing the formulation of appropriate public health policies. However, surveillance could be made more effective. We have determined that the time taken from strain collection to genome submission for over 1.7 million SARS-CoV-2 strains available at GISAID. We find that strain-wise, time lag in this process ranges from one day to over a year. Country-wise, the UK has taken a median of 16 days (for 417,287 genomes), India took 57 days (for 15,614 genomes), whereas Qatar spent 289 days (for 2298 genomes). We strongly emphasize that along with increasing the number of genomes of COVID-19 positive cases sequenced, their accelerated submission to GISAID should also be strongly encouraged and facilitated. This will enable researchers across the globe to track the spreading of variants in a timely manner; analyse their biology, epidemiology, and re-emerging infections; and define effective public health policies.

## Introduction

Genomic surveillance of the evolving SARS-CoV-2 strains is an important tool to help control the raging pandemic^1^. For efficient surveillance, the first major requirement is the availability of all sequenced genomes on an open-access platform that is accessible by researchers worldwide, to enable them to analyze how this virus is evolving and spreading. Therefore, soon after researchers became aware of COVID-19, towards the end of 2019, they started depositing the sequenced genomes to the Global Initiative on Sharing All Influenza Data (GISAID), a pre-existing platform for all influenza viruses. As of now, GISAID is the largest open-access portal, hosting the genome sequence and related epidemiological and clinical data of more than 1.7 million SARS-CoV-2 strains. In a mere 1.5 years, this virus has become one of the most studied organisms in history, with GISAID playing a major enabling role. Thanks to ongoing genomic surveillance using this data, several new variants such as B.1.1.7 (Alpha; first in the UK); B.1.351 (Beta; first in South Africa); B.1.1.28 (Gamma; P.1, first in Brazil); B.1.617.2 (Delta; first in India); B.1.617.1 (Kappa; first in India); P.3 (Theta; first in the Philippines); and B.1.427 and B.1.429 (Epsilon; first in the USA) have been identified^2–5^. This information has been used worldwide to update public health policies to control COVID-19 infections^6,7^.

Considering the benefits of genomic surveillance^6,8^, scientists have pressured countries to increase their sequencing capacity, and this has led to several initiatives such as <u>COG-UK</u>, <u>INSA-COG</u> (India), <u>NGS-SA</u> (South Africa), <u>SPHERES</u> (USA), etc. However, besides increasing the fraction of samples sequenced, there is another issue that scientists need to be concerned about i.e. “How soon are the sequences being submitted to GISAID or any other open access platform?” Rapid submission will enable the international community to analyse the variants emerging around the world quickly and provide actionable information to governments.

## Methods

Uisng the lastest data (as of 27 May 2021) available at GISAID, we have calculated the Collection to Submission Time Lag (CSTlag) per strain. We have also calculated the median and average CSTlag time for each country and continent (category 1: for all countries and category 2: for all those countries who have submitted over 1000 genomes). Country population and total COVID-19 cases data were obtained from Worldometer on June 02, 2021, 17:32 GMT. Based on these information, we have also calculated the rate of genome sequencing normalized with total COVID-19 cases and one million population per country respectively.

## Results

Our statistical analysis (Figure 1 and S1; Table S1 and S2) for 1,718,035 SARS-CoV-2 strains submitted to GISAID has determined that the Collection to Submission Time Lag (CSTlag) per strain ranges from 1 day to over a year. Examining the median CSTlag values for countries that have sequenced >1000 SARS-CoV-2 genomes, we note that the CSTlag from the UK is the shortest i.e., 16 days for over 417,000 genomes. For the rest of Europe, the lag is 25 days for over 590,000 genomes. The USA has spent almost 26 days for over 498,000 genomes, whereas for Canada it is 88 days for over 44,000 genomes. Amongst the Oceania countries, the CSTlag for New Zealand is 40 days for over 1000 genomes, whereas for Australia, it is 51 days for over 17,000 genomes. In Asia, the median lag is 72 days, for over 89,000 genomes, with Singapore having the shortest lag of 26 days for 2405 genomes, and Qatar the longest lag of 289 days for 2298 genomes. India’s CSTlag is 57 days for 15,614 genomes whereas Japan, which has sequenced the most genomes in Asia, has taken 79 days for over 37,000 genomes. For South American countries, the median lag is 61 days for over 18,000 genomes, whereas countries in Africa have taken 50 days for over 7000 genomes (Table S2).

**Figure 1:**
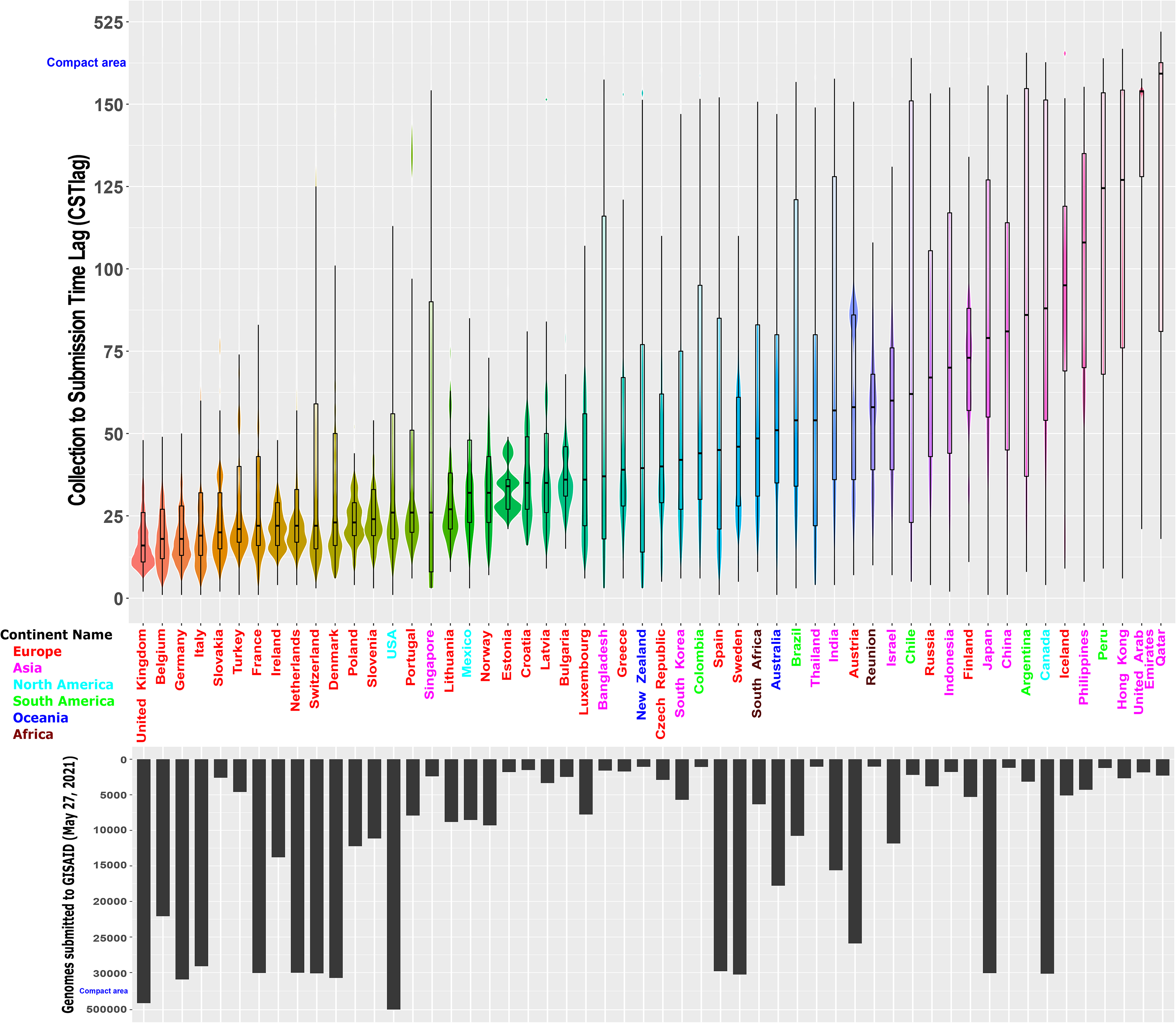
Violin plot illustrating the CSTlag values for the 54 countries that have sequenced over 1000 genomes. The box plot inside the violin plot depicts the median CSTlag per country. Outlier CSTlag entries per country are not shown in this illustration. Each country’s name is color-coded according to the continent. We have also mapped the relative distribution of the number of genome sequences submitted by each country as a bar plot.

Coming to the rate of sequencing, top-performing countries Iceland, Australia, New Zealand, and Denmark have sequenced approximately 77%, 59%, 39%, and 35% of their positive cases respectively (Table S1). The USA and UK have sequenced over 400,000 genomes each, which is 9.2% and 1.5% of their respective positive cases. India, being the second-largest country based on both total population and known COVID-19 cases, has sequenced a mere 0.05% of the reported cases. On average, African, Asian, and South American countries have sequenced a mere 0.36%, 0.21%, and 0.07% of their total COVID-19 cases, whereas this number is 1.9%, 1.4%, and 37% of European, North American, and Oceania countries. Population-wise, most of the European countries, the USA, Israel (Asia) and Reunion (Africa) have sequenced samples from over 1000 people per million population (ppmp). Amongst countries with over 100 million population, including Brazil (50 ppmp), India (11 ppmp), Indonesia (6 ppmp), Nigeria (4 ppmp), and Pakistan (1 ppmp), only the USA (1,497 ppmp) and Japan (297 ppmp) have sequenced over 100 ppmp. Cumulatively, African, Asian, and South American countries have carried out the genome sequencing of only 14, 21, and 49 ppmp, whereas this number is 1198, 948, and 607 ppmp for European, North American, and Oceania countries (Table S2).

## Discussion

In terms of the delay in sequence submissions, there may be several reasons for this. The speed of sequence submission to GISAID is based on (i) the time taken from sample collection from a COVID-19 patient to RNA isolation in the lab and its dispatch to the sequencing centre and (ii) the time from RNA sample arrival at the sequencing centre to the uploading of the sequence. Countries with a shorter median CSTlag are more likely to have strong public health systems allowing efficient sample and metadata collection, and smoother coordination between the sample collection centre, the RNA isolation lab, and the sequencing lab. Countries without such a strong system would be at a disadvantage and may face additional logistical problems in sample/metadata collection and shipping because of lockdown-related restrictions. Several countries might have a shortage of labs that can handle COVID-19 samples, or might have an overly centralized system, wherein only a few labs are authorized to handle such samples, causing a delay in sequencing and submission. A paucity of funds or restrictions on importing reagents and equipments would also add to the delay. The use of older sequencing technologies that are low-throughput and more expensive per sample would complicate matters further.

Most of the countries with a short CSTlag are industrialized nations that are likely to have strong linkages between the clinical and scientific establishments. This is not always so for other countries. Many countries with a longer CSTlag have a less developed public health system. They might have had to establish novel collaborations and institutional arrangements to help deal with the pandemic. All of this would have taken time, which would have impacted work on the ground. Some of the possible causes for delay listed above are known to have been true in India, for instance, and are being resolved^9,10^.

Sometimes, even after rapid sequencing, genomes may not be promptly uploaded to GISAID, and there may be several reasons for that. First, the importance of genomic surveillance may not have been well understood, especially in the early months of the pandemic. Second, there may be a wish to withhold information, in order to publish or patent first. Third, several governments may be sensitive to the issue of virulent strains, in particular, being named after their countries. The WHO initiative of renaming variants with Greek letters may help in resolving this issue^5^. Finally, in many countries, there may be significant bureaucratization or political interference at various steps from sample collection to uploading sequences to GISAID, which adds to the delay. Although one does not know the extent of various problems in each country, it is likely that far more samples have been sequenced than are represented in GISAID.

In countries with a longer CSTlag, the sequenced variants may have enough time to establish themselves across the region, or – based on a significant mutation rate^11^ – may evolve into completely new variants, if quick tracking, tracing, and actions to stop transmission are not undertaken. Therefore, this issue must receive urgent attention. All bottlenecks that prevent a lower CSTlag must be addressed.

Overall, an effective genomic surveillance system requires not only sequencing a significant fraction of strains from COVID-19 patients, but also rapid genome submission to open access platforms like GISAID. This will enable researchers across the globe to track the evolved variants, their mutations, epidemiology, and biological consequences, which will provide crucial inputs for appropriate and effective public health policies.

## Supplementary Data

**Table S1:** The population of each country and the country-wise distribution of (i) total COVID-19 cases, (ii) genomes submitted to GISAID, (iii) rate of genome sequencing normalized with COVID-19 cases, (iv) rate of genome sequencing normalized with one million population (v) average CSTlag and (vi) median CSTlag values.

**Table S2: Continent-wise statistics.** (i) Total population of the respective continent, (ii) Total COVID-19 cases reported in the respective continent, (iii) Total genomes submitted to GISAID from the respective continent, (iv) rate of genome sequencing in the continent normalized with COVID-19 cases, (v) rate of genome sequencing in the continent normalized with one million population, (vi) average CSTlag for all strains per continent, and (vii) median CSTlag values for all strains per continent.

**Figure S1**: Violin plot illustrating the CSTlag values for the 54 countries that have sequenced over 1000 genomes. The box plot inside the violin plot depicts the median CSTlag per country. All CSTlag entries per country are shown in this illustration. Each country’s name is color-coded according to the continent. We have also mapped the relative distribution of the number of genome sequences submitted by each country as a bar plot.

## Views

The views expressed in this letter are those of the authors, and not necessarily those of either funding agency or any other institution.

