## Supplementary figures and images for "Country-wide genomic surveillance of SARS-CoV-2 strains"

### Figure S1---With outliers.pdf

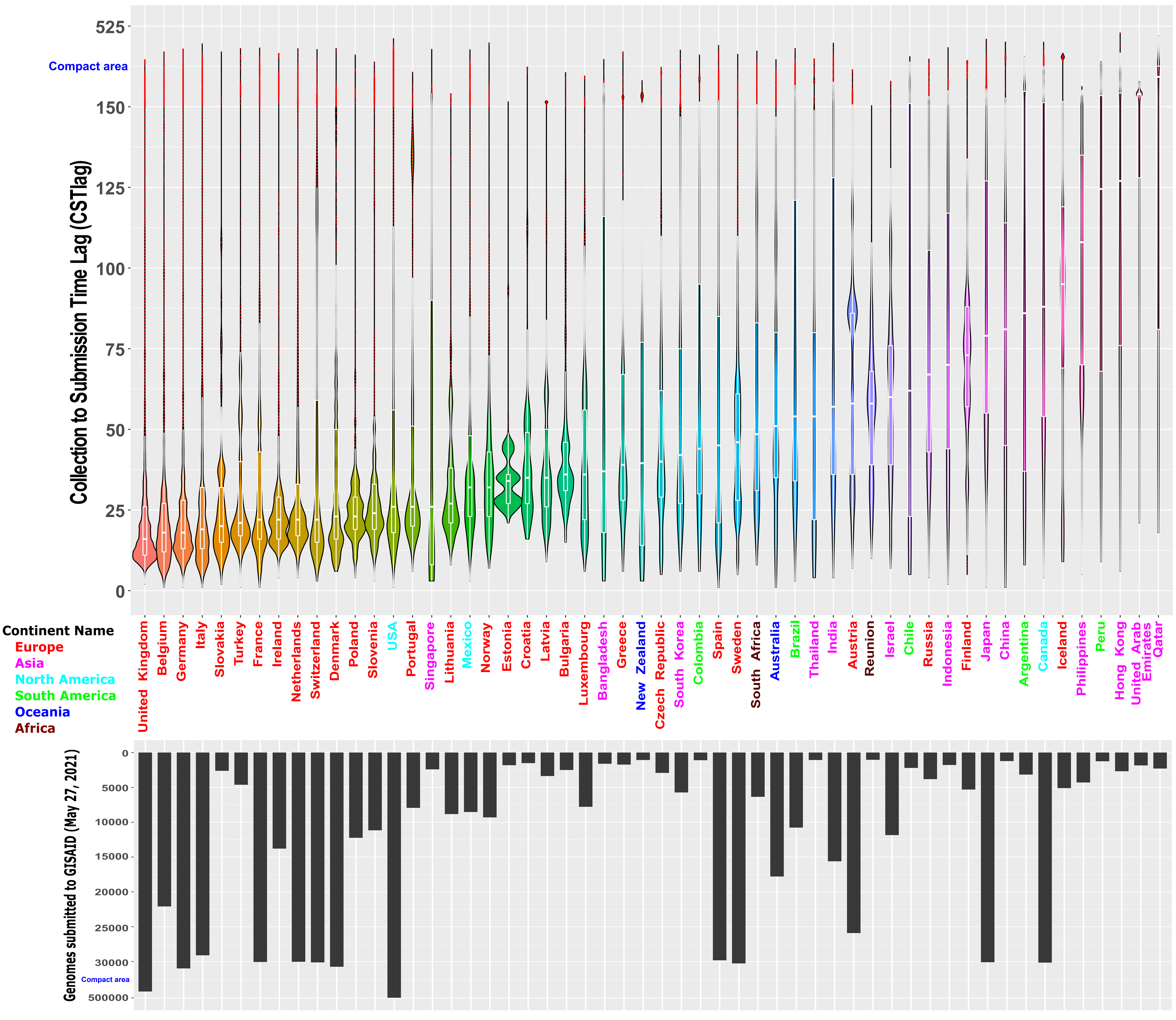
